# A two-component system signaling hub controls enterococcal membrane remodeling in response to daptomycin

**DOI:** 10.1101/2025.11.16.688641

**Authors:** Cristina Colomer-Winter, Zeus J. Nair, Jerome Y. J. Chua, Soukayna Jabli, Mélanie Roch, Amaury Cazenave-Gassiot, Roberto Sierra, Diego O. Andrey, Shu-Sin Chng, Kimberly A. Kline

**Affiliations:** Department of Microbiology and Molecular Medicine, University of Geneva, Geneva, 1211 Switzerland; Singapore-MIT Alliance for Research and Technology, Antimicrobial Drug Resistance Interdisciplinary Research Group, Singapore, 138602 Singapore; Singapore Centre for Environmental Life Sciences Engineering, Nanyang Technological University, 637551 Singapore; Singapore Lipidomics Incubator, Life Sciences Institute, National University of Singapore, 117456 Singapore; Department of Biochemistry, Yong Loo Lin School of Medicine, National University of Singapore, 117596 Singapore; Division of Infectious Diseases, Department of Medicine, Geneva University Hospitals and Medical School, Geneva, 1211 Switzerland; Division of Laboratory Medicine, Department of Diagnostics, Geneva University Hospitals and University of Geneva, Geneva, 1211 Switzerland; Department of Chemistry, National University of Singapore, 117543 Singapore

**Keywords:** Two-component systems, daptomycin resistance, *Enterococcus faecalis*, LTA, lipid remodeling

## Abstract

Daptomycin is a last resort antibiotic used to treat vancomycin-resistant enterococcal infections, but daptomycin resistance (DAP^R^) arises quickly during treatment. Resistance is due to sequential acquisition of point mutations in the two-component system LiaFSR and in cardiolipin synthases and is associated with alteration of phospholipid and glycolipid membrane composition. The molecular mechanisms underlying these lipid changes are currently unknown. Similarly, it is unclear why mutations in *liaFSR* occur prior to mutations in *cls*. We discovered that *Enterococcus faecalis* remodels membrane composition as a phenotypic response to daptomycin that parallels the membrane composition of DAP^R^ strains. The enrichment in glycolipids that follows antibiotic exposure is due to LtaS1, the main LTA synthase of *E. faecalis*. Moreover, LtaS1 activity is governed by a network of two-component systems formed by LiaFSR, SapRS, and BsrRS that couple antibiotic sensing with membrane lipid remodeling. Together, our results provide a unifying mechanism that drives phenotypic membrane fortification in a Gram-positive pathogen which simultaneously predisposes the cell to acquire genetic high-level daptomycin resistance.

**Significance Statement:** Daptomycin is the preferred alternative to treat vancomycin-resistant *Enterococcus* infections. However, the efficacy of daptomycin is limited by the acquisition of daptomycin resistance. Alterations in lipid membrane composition represent a conserved strategy to fortify the Gram-positive membrane. We discovered that *E. faecalis* phenotypically remodels membrane composition in response to daptomycin and that the enrichment in glycolipids is dependent on LTA synthesis. We expand our limited knowledge of enterococcal LTA synthases, confirming LtaS1 as the main LTA synthase and placing LTA biogenesis under control of a network of antibiotic-responsive two-component systems. Our work provides a molecular explanation for the sequential evolution of daptomycin resistance and may support the use of existing LtaS inhibitors to prevent acquisition of daptomycin resistance.

## Introduction

*Enterococcus faecalis* and *Enterococcus faecium* are Gram-positive commensals of the human gastrointestinal tract. However, Enterococci are also leading pathogens of hospital-associated infections such as bloodstream and catheter-associated urinary tract infections (1). Due to the multidrug resistance of clinical strains, treatment is often difficult and results in ∼450,000 annual deaths worldwide (2). Vancomycin-resistant Enterococci (VRE) are especially concerning; they are classified as ESKAPE pathogens and are considered a global threat to public health. Thus, a mechanistic understanding of antimicrobial resistance is urgently needed to improve current anti-infective strategies.

VRE infections can be treated with daptomycin (DAP), a last resort lipopeptide antibiotic (1). Similar to cationic antimicrobial peptides (CAMPs), it displays a dual mode of action: it depolarizes the cell membrane by binding to the anionic phospholipid phosphatidylglycerol (PG), and it inhibits cell wall synthesis through tripartite interaction with PG and the peptidoglycan precursor lipid II at the cell septum (3–5). Though DAP is initially potent, daptomycin resistance (DAP^R^) arises quickly. In *E. faecalis*, DAP^R^ occurs in two consecutive steps: first cells acquire mutations that constitutively activate the two-component system (TCS) LiaFSR, and then cells acquire secondary mutations in phospholipid biosynthetic genes that drive antibiotic resistance beyond the clinical breakpoint (6–8). The enterococcal Lia system is composed of a histidine kinase (HK, LiaS), a response regulator (RR, LiaR), and a set of accessory proteins (LiaF, LiaX, LiaY, LiaZ) (9). The current model is that activation of LiaFSR promotes resistance through two mechanisms: it activates transcription of CAMP resistance factors, and it displaces cardiolipin synthases (Cls) away from the cell septum (9–12). The latter leads to spatial rearrangement of anionic lipid microdomains that attract DAP to non-septal locations, thus protecting the septum – the cellular nexus for peptidoglycan synthesis and cell division – from antibiotic damage (4, 5, 9).

Less is known about secondary mutations in phospholipid biosynthetic genes that confer high-level DAP^R^ to Enterococci. In other Gram-positive bacteria, mutations that decrease the concentration of PG in the membrane lead to DAP^R^ (13, 14). Resistance stems from the fact that PG is the primary target of DAP and is required for membrane depolarization and peptidoglycan synthesis inhibition. DAP^R^ *E. faecalis* strains also have lower PG levels (14–17). The most frequent secondary mutation is a gain-of-function of Cls1, a cardiolipin synthase that uses PG as a substrate to synthetize the phospholipid cardiolipin (CL) (18). Thus, secondary mutations are thought to confer enterococcal high-level DAP^R^ because they decrease PG levels (17). However, this hypothesis is challenged by deletion of phospholipid biosynthetic genes that lower membrane PG but do not confer DAP^R^ in *E. faecalis* (19, 20). Moreover, it is still unknown why secondary mutations such as in *cls1* are only acquired after constitutive activation of LiaFSR (8). These findings illustrate a gap in our understanding of how *E. faecalis* modulates membrane composition in response to antibiotics.

Here, we discovered that *E. faecalis* remodels its membrane composition as part of a phenotypic adaptive response to DAP. Exposure to DAP decreases PG levels and in turn increases the glycolipid diglucosyl diacylglycerol (DGDAG). Moreover, DAP triggers recycling of the phospholipid lysyl-phosphatidylglycerol (LPG) and induces lipoteichoic acid (LTA) synthesis. Applying a combination of transcriptomic, lipidomic, and genetic analyses, we identified lipoteichoic acid synthase 1 (LtaS1) as the key enzyme responsible for *de novo* glycolipid synthesis during antibiotic stress. LTA production is under the control of a TCS network formed by LiaFSR, SapRS, and BsrRS which coordinate the antibiotic stress response. Our findings reveal how *E. faecalis* responds to DAP and suggest a mechanism by which LiaFSR-dependent membrane remodeling promotes acquisition of secondary mutations in DAP^R^ strains.

## Results

### DAP triggers adaptive lipid remodeling in *E. faecalis* OG1RF

In *E. faecalis,* DAP^R^ arises due to a primary mutation in LiaFSR followed by a secondary mutation in phospholipid biosynthetic genes (8), suggesting that LiaFSR alters the membrane to become permissive to secondary mutations. Since DAP triggers LiaFSR activation in DAP-susceptible (DAP^S^) strains (10), we hypothesized that *E. faecalis* might transiently remodel its membrane composition as an adaptive response to the antibiotic. To determine if exposure to DAP triggers lipid remodeling, we incubated exponentially grown *E. faecalis* OG1RF with a sub-inhibitory concentration of DAP (2 µg/mL) (**Fig. S1A**) prior to lipid analysis by liquid chromatography-tandem mass spectrometry (LC-MS/MS). We quantified total membrane content of PG, LPG, and DGDAG following DAP treatment since DAP^R^ *E. faecalis* membranes contain less PG and LPG and more DGDAG than DAP^S^ strains (14–16). Compared to untreated cells, DAP induced a significant decrease in PG and LPG, accompanied by a notable yet not statistically significant increase in DGDAG, fully recapitulating the trends observed in DAP^R^ strains (**Fig.1A**). We also quantified individual PG, LPG, and DGDAG lipid species which are defined by the carbon length and saturation of their fatty acids. DAP triggered a decrease in PG and LPG phospholipid species of 32 to 34 carbons in length and a significant increase in DGDAG glycolipid species (**Fig. S1B)**. Fatty acids detected in the three lipid classes, including the predominant species 34:1, were enriched in DGDAG to the detriment of PG and LPG (**Fig. 1B**). These results confirmed that *E. faecalis* remodels its membrane composition as a rapid response to DAP.

**Figure 1.**
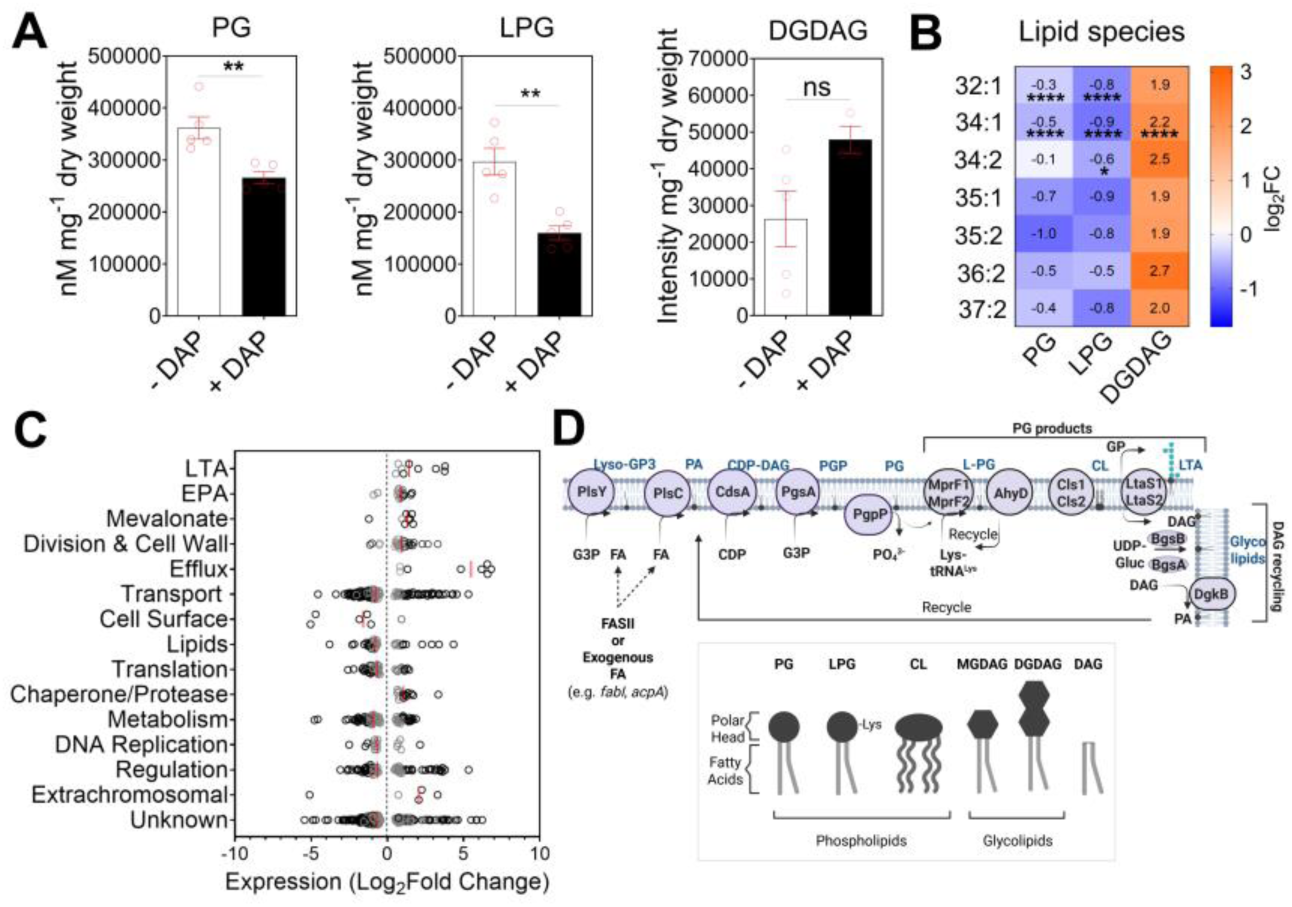
Lipidomic and transcriptomic changes in *E. faecalis* OG1RF upon DAP treatment. (A) LC-MS/MS quantification of total PG, LPG, and DGDAG membrane content of treated and untreated *E. faecalis* OG1RF. Exponentially grown cells were treated with 2µg/mL DAP for 1 hour prior to lipid extraction. PG and LPG were quantified and normalized to internal standard and cell pellet weight, while DGDAG intensity was normalized to cell pellet weight for a semi-quantitative measurement. Each bar represents the mean ± standard error of 3 to 5 biological replicates. Differences in lipid content were analysed via Student’s t test. (** P ≤ 0.01). (B) Heat map summarizing log_2_-fold changes in PG, LPG, and DGDAG lipid species upon DAP treatment. Fatty acids common to the three lipid classes are shown. Statistically significant changes are labelled (* P ≤ 0.05, **** P ≤ 0.0001). (C) RNA-Seq of DAP-treated OG1RF. Exponentially grown cells were treated with 2µg/mL DAP for 15 minutes prior to RNA extraction. Analysis was performed on 3 biological replicates per condition. The graph shows the functional classification of differentially expressed genes. Genes exhibiting Log_2_FC ≥ 1.0 with FDR ≤ 0.05 when compared to untreated OG1RF are shown in black. (C) Schematic of phospholipid biosynthesis and recycling in Enterococci. G3P: Glycerol-3-phosphate, FA: Fatty Acids, FASII: Endogenous FA biosynthesis pathway, PA: Phosphatidic acid, CDP-DAG: CDP-diacylglycerol, PGP: Phosphatidylglycerol phosphate, PG: Phosphatidylglycerol, LPG: Lysyl-phosphatidylglycerol, CL: Cardiolipin, DAG: Diacylclycerol, UDP-Gluc: UDP-Glucose. MGDAG: Monoglucosyl diacylglycerol, DGDAG: Diglucosyl diacylglycerol. Figure created with Biorender.

### DAP alters transcription of fatty acid and lipid biosynthetic pathways

To determine if regulation of adaptive membrane remodelling occurred on the transcriptional level, we performed RNA-Seq on DAP-treated *E. faecalis* OG1RF. Sub-inhibitory DAP elicited 683 differentially expressed genes (FDR < 0.05) compared to untreated controls (**Fig. 1C, Dataset S1**). Consistent with previous studies (10, 21), transcription of *liaFSR* and *liaXYZ* was strongly induced. LiaFSR activates transcription of *sapRS*, the bacitracin-sensing TCS of *E. faecalis* (also known as YxdJK or MadRS) (11, 12, 22). Accordingly, SapRS-regulated genes such as *mprF2, dltABCD,* and *madEFG* governing CAMP resistance (12, 23, 24) were highly expressed in response to DAP. We hypothesized that transcriptional repression of PG biosynthesis could lower membrane PG levels. Most genes of the PG biosynthetic pathway (*plsY*, *plsC*, *cdsA*, *pgsA*, and *pgpP* (**Fig. 1D**) (25)) were differentially expressed in response to DAP, although without uniform directionality (**Fig. S1C,Dataset S1**). A clearer trend appeared in the fatty acid biosynthetic pathway (FAS-II) that feeds into phospholipid synthesis. DAP triggered decreased expression of almost the entire pathway, including rate limiting enzymes *fabI* and *acpA* (26, 27), suggesting that the decrease in total PG levels could be partially due to transcriptional repression of fatty acid biosynthesis. Alternatively, PG content can be decreased by induction of enzymes that consume PG as a substrate. PG is a common precursor of all other phospholipids and glycolipids, providing fatty acids on which the polar head of the respective lipid class will be attached (**Fig. 1D**). PG-consuming enzymes in Gram-positive bacteria include Cls (CL synthesis), MprF (LPG synthesis), and LtaS (LTA synthesis) (28). In DAP-treated *E. faecalis,* transcription of *cls1* was unaffected, while *cls2* was slightly decreased. However, *mprF1*, *mprF2,* and two predicted *ltaS* genes were induced, suggesting higher PG consumption by LPG and LTA synthases **(Fig. S1C, Fig. S1D)**.

### Phospholipids are recycled into glycolipids and LTA in response to DAP

Lipidomic quantification showed a decrease in PG and LPG, and an increase in DGDAG following DAP exposure. Transcriptomic analysis suggested that the decrease in PG was due to both lower PG synthesis and higher PG consumption by MprF and LtaS. All LTA synthases (LtaS) use PG to synthesize LTA and DGDAG (**Fig. 2A**) (29). We hypothesized that the alteration in phospholipid (PG and LPG) and glycolipid (DGDAG) content was because newly synthesized PG was primarily consumed for DGDAG production. Alternatively, but non-mutually exclusive, it could be that pre-existing PG and LPG are recycled into DGDAG since these lipids are recycled into PG in other Gram-positive organisms (**Fig. 1D**, **Fig 2A)** (30–32).

**Figure 2.**
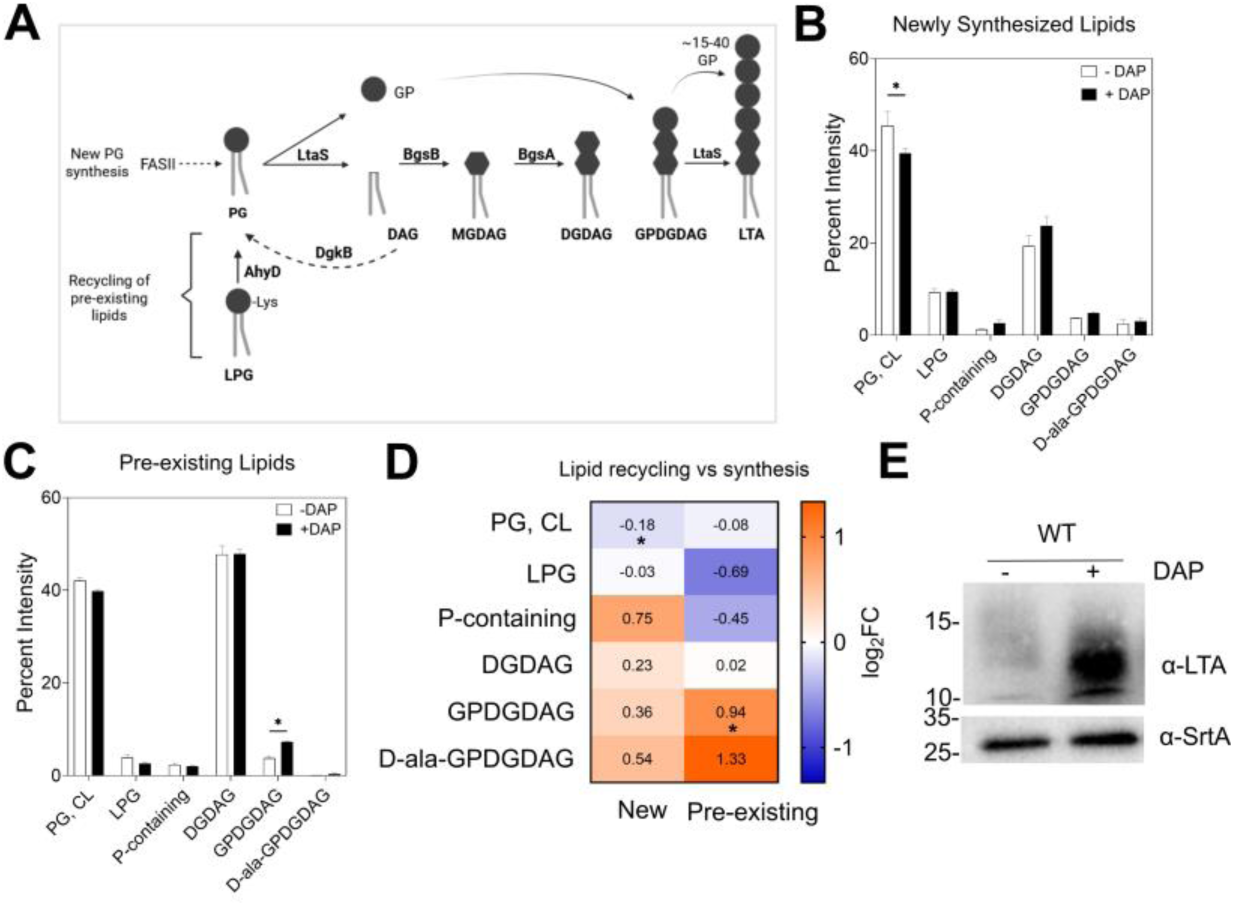
DAP triggers *de novo* glycolipid synthesis and recycling of phospholipids into glycolipids and LTA. (A) Schematic of glycolipid and LTA synthesis. LtaS cleaves PG into glycerol phosphate (GP) and DAG. DAG is utilized by glycosyltransferases BgsB and BgsA to produce the glycolipids MGDAG and DGDAG, respectively. In parallel, LtaS polymerizes 15 to 50 GP subunits onto a DGDAG glycolipid which anchors the LTA structure to the membrane. There are two sources of PG: *de novo* PG synthesis from newly synthetized fatty acids (FAS-II) and glycerol-3-phosphate, or recycled PG from DAG and LPG by AhyD and DgkB, respectively. Figure created with Biorender. (B, C) Densitometric analysis of [^14^C]-radiolabelled 1D TLC spots of (B) newly synthesized lipids and (C) pre-existing lipids in response to DAP, displaying the percentage intensities of each spot relative to the total intensity of all lipid classes detected. For newly synthesized lipids, [^14^C]-acetate was added during DAP exposure, while for pre-existing lipids, [^14^C]-acetate was added at the start of the sub-culture followed by addition of unlabeled acetate during DAP exposure. Each bar represents the mean ± standard error of measurement calculated from 3 biological replicates. Spot identities were determined using lipid standards. Differences were analyzed via two-way ANOVA followed by Šídák’s post-test (* P ≤ 0.05). (D) Heat map summarizing log_2_-fold changes in new and pre-existing PG, LPG, and DGDAG lipid classes after DAP treatment. Statistically significant changes are labelled (* P ≤ 0.05). (E) LTA Western Blot of treated and untreated cell extracts. Sortase A (SrtA) was used as a loading control. Representative blot of three independent biological replicates. Size is denoted in kDa.

To distinguish newly synthesized lipids (following DAP treatment) from pre-existing lipids (present before DAP exposure), we performed pulse-chase thin layer chromatography (TLC) of [14C]-radiolabeled lipids. TLC allows for detection of additional lipidic entities, including a previously reported P-containing lipid of unknown identity and glycolipids committed to LTA assembly – glycerophospho-diglucosyl-diacylglycerol and it’s D-alanine modified form (GPDGDAG, D-Ala-GPDGDAG) (19). CL resolves at the same spot as PG and the two lipids cannot be distinguished under these conditions. When exposed to DAP, we detected a drop in newly synthesized PG and CL and an accumulation of newly synthesized DGDAG (**Fig. 2B**). By contrast, pre-existing LPG modestly decreased and GPDGDAG, the precursor of LTA, increased (**Fig. 2C**) (16). Log^2^-fold changes between treated and untreated cells highlighted these findings (**Fig. 2D**), collectively indicating that newly synthesized PG was rapidly consumed to produce DGDAG and that pre-existing phospholipids such as LPG were recycled into the DGDAG anchor of LTA. The results imply that DAP triggers the cell to broadly redirect all available phospholipids towards LTA production (33). Indeed, immunoblotting confirmed increased LTA synthesis following DAP treatment (**Fig. 2E**).

### LtaS1 is responsible for LTA production in *E. faecalis*

*E. faecalis* is predicted to have two LtaS enzymes based on protein homology to other Gram-positive LtaS, but their activity had not been experimentally confirmed (29). We named them *ltaS1* (OG1RF_11033) and *ltaS2* (OG1RF_11521) based on chromosomal position. Transcriptional induction of *ltaS1* and *ltaS2* after DAP exposure suggested that glycolipid remodelling and increased LTA production was due to the activity of at least one of them. We generated single (Δ*ltaS1*, Δ*ltaS2*) and double Δ*ltaS1*Δ*ltaS2* deletion strains. Whole genome sequencing confirmed the absence of additional mutations in Δ*ltaS2* and Δ*ltaS1*Δ*ltaS2* strains, while Δ*ltaS1* harboured a SNP in a gene of unknown function (OG1RF_RS07860). LTA immunoblots showed that neither Δ*ltaS1* nor Δ*ltaS1*Δ*ltaS2* produced detectable LTA, even when exposed to DAP, whereas Δ*ltaS2* retained the ability to synthesize LTA (**Fig. 3A**). Moreover, Δ*ltaS1* was 16 times more sensitive to DAP than wild-type (WT) and Δ*ltaS2* (**Fig. 3B**). Complementation of *ltaS1* restored LTA production and DAP susceptibility to WT levels (**Fig. S2A, Fig. 3C**). These data indicate that LtaS1 is the main LTA synthase of *E. faecalis* and that it contributes to baseline DAP sensitivity.

**Figure 3.**
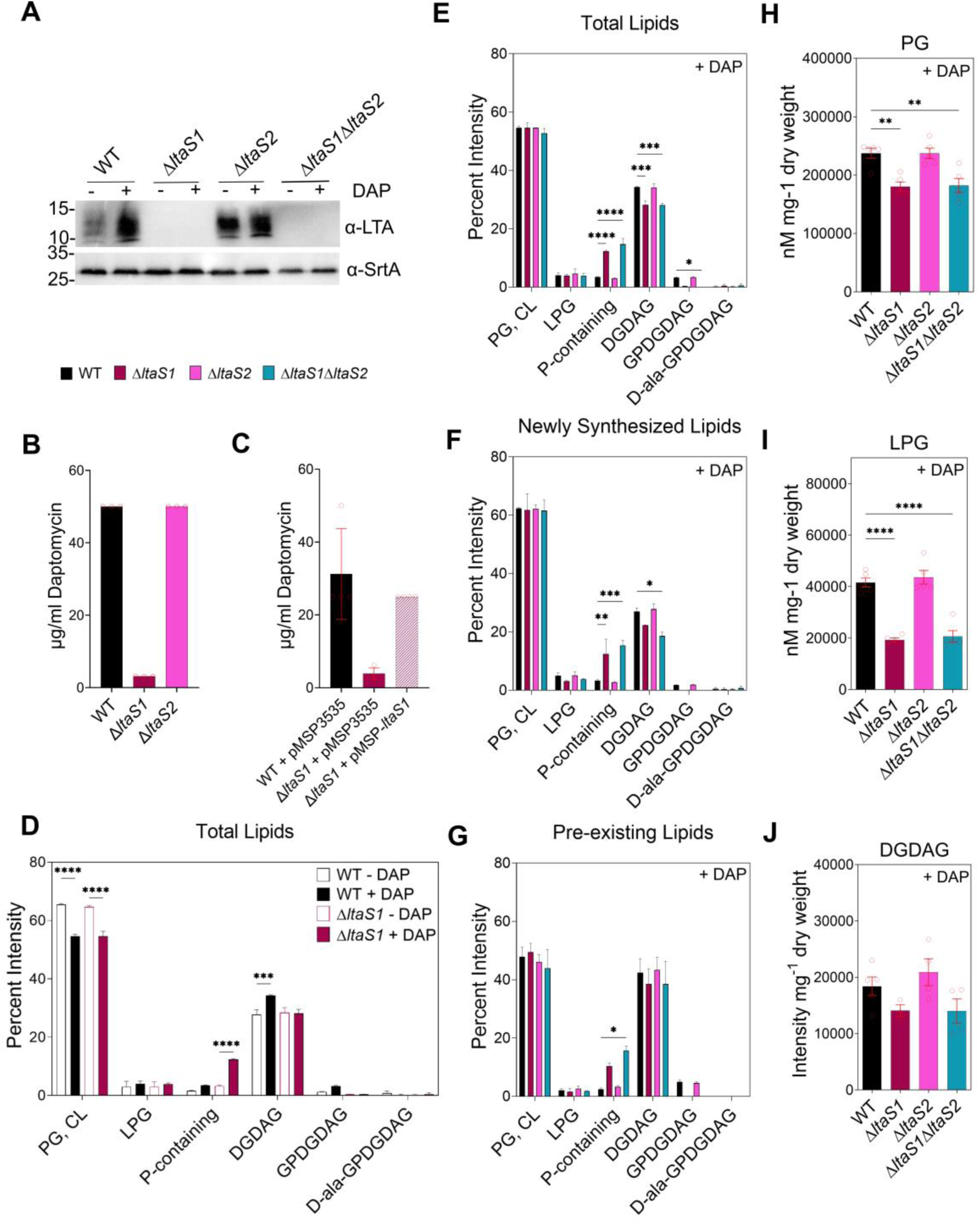
LTA synthetized by LtaS1 drives glycolipid membrane remodelling. (A) LTA Western Blot of untreated and DAP-treated OG1RF WT, Δ*ltaS1*, Δ*ltaS2*, and Δ*ltaS1*Δ*ltaS2* cell extracts. SrtA was used as a loading control. Representative blot of two independent biological replicates. Size is denoted in kDa. (B) DAP MIC of OG1RF WT, Δ*ltaS1* and Δ*ltaS2* strains in Ca-adjusted BHI media. A final inoculum of ∼10^5^ CFU/mL was challenged with increasing concentrations of DAP. After 18-20h incubation at 37°C, MIC was determined by visually inspecting for the lowest DAP concentration that completely inhibited bacterial growth. (C) DAP MIC of OG1RF WT, Δ*ltaS1* and the complemented Δ*ltaS1* strains in Ca-adjusted BHI media. The expression vector pMSP3535 was used to complement *ltaS1* in *trans* (pMSP-*ltaS1*). (D, E, F, G) DAP-dependent lipid remodelling analysed by TLC. Densitometric analysis of [^14^C] radiolabeled 1D TLC spots of (D, E) total lipids, (F) newly synthetized lipids, and (G) pre-existing lipids in parent and LTA-defective strains after DAP treatment (+DAP). For visualization purposes, graph (D) shows a side-by-side comparison of total lipid content between the parent and Δ*ltaS1* strains under treated (+DAP) and untreated (-DAP) conditions. Graphs in (E, F, G) show comparisons between the entire panel of strains. Each bar represents the mean ± standard error of measurement calculated from 2 biological replicates. Spot identities were determined using lipid standards. Differences were analyzed via a two-way ANOVA (* P ≤ 0.05, ** P ≤ 0.01, *** P ≤ 0.001, **** P ≤ 0.0001), (H, I, J) LC-MS/MS quantification of total (H) PG, (I) LPG, and (J) DGDAG content of treated WT and LTA-deficient *E. faecalis* cells. PG and LPG were normalized to internal standard and cell pellet weight, while DGDAG intensity was normalized to cell pellet weight for a semi-quantitative measurement. Each bar represents the mean ± standard error of 3 to 5 biological replicates. Differences in lipid content were analysed via one-way ANOVA. (** P ≤ 0.01, **** P ≤ 0.0001).

### LtaS1 remodels glycolipid membrane composition in response to DAP

We next asked whether LtaS1 was responsible for changes in phospholipid and glycolipid content following DAP exposure. Deletion of *ltaS1,* either alone or in combination with *ltaS2,* abolished the DAP-induced increase in DGDAG (**Fig. 3D**, **Fig. 3E**), consistent with the role of LtaS enzymes in glycolipid synthesis in other organisms (29). Membranes of Δ*ltaS1* and Δ*ltaS1*Δ*ltaS2* contained PG and CL comparable to WT but had ∼4 times more P-containing lipid. These alterations were DAP-specific, as no major differences in lipid content were observed in the absence of DAP (**Fig. S2B**). Consistent with LTA immunoblots, GPDGDAG levels were reduced in the Δ*ltaS1* mutant and undetectable in the Δ*ltaS1*Δ*ltaS2* strain. The membrane composition of Δ*ltaS2* was similar to the parent strain across lipid classes, collectively suggesting that LtaS2 may provide auxiliary- or primase-like activity similar to LtaP in *Listeria monocytogenes* (29).

Next, we traced *de novo* lipid synthesis and recycling to determine if the LtaS1-dependent increase in DGDAG was fuelled by new or pre-existing lipids. Following DAP exposure, Δ*ltaS1* and Δ*ltaS1*Δ*ltaS2* strains failed to accumulate newly synthesized DGDAG (**Fig. 3F**). Moreover, deletion of *ltaS1* led to a significant accumulation of both newly synthesized and pre-existing P-containing lipid (**Fig. 3G**). No significant differences were observed in the absence of DAP (**Fig. S2C-D**). These results confirmed that LtaS1 was required for *de novo* synthesis of DGDAG in response to DAP. Next, we sought to determine if LtaS1 was driving the decrease in PG due to its higher demand for LTA. We used LC-MS/MS to distinguish PG from CL. Unexpectedly, PG was even lower in Δ*ltaS1* and Δ*ltaS1*Δ*ltaS2* strains after antibiotic treatment (**Fig. 3H**). We quantified LPG and DGDAG in *ltaS*-deficient strains to confirm our TLC results. Deletion of *ltaS1* led to significantly lower LPG even in the absence of DAP (**Fig. 3I, Fig. S2F**), and abolished accumulation of DGDAG in response to DAP (**Fig. 3H**). Collectively, lipidomic profiling confirmed that LtaS1 drives adaptive glycolipid remodelling in response to DAP.

### A TCS network controls LTA synthesis in response to DAP

Our results demonstrated that *E. faecalis* mounts a protective response to DAP by increasing LTA and glycolipid synthesis in a LtaS1-dependent manner. Since *ltaS1* transcription was induced in response to DAP and LiaFSR regulates the DAP stress response, we hypothesized that *ltaS1* could be under the control of LiaFSR. We therefore defined the LiaFSR regulon. As previously reported (10), deletion of *liaFSR* under unstressed conditions led to differential expression of 16 genes (**Fig. S3A**, **Dataset S2**). When exposed to DAP, we observed an additional 72 differentially expressed genes, including LiaFSR- and SapRS-dependent CAMP resistance factors such as *mprF2*, *dltABCD*, and *madEFG* (**Fig. 4A, Dataset S2**) (10–12). Moreover, transcription of several genes involved in lipid function was altered, including the flotillin *floT* involved in membrane microdomain organization (34), and *gdpD,* a glycerophosphoryldiester phosphodiesterase previously associated with DAP^R^ in *E. faecalis* (6). The *ltaS1* and *ltaS2* genes were not differentially expressed; however, expression of the DGDAG synthase *bgsA* was lower in the Δ*liaFSR* than in the parent strain. Transcriptomic analysis of a Δ*liaR* strain under the same conditions yielded comparable results (**Fig. S3B, S3C, Dataset S2**), suggesting an insulation of the LiaFSR signaling cascade.

**Figure 4.**
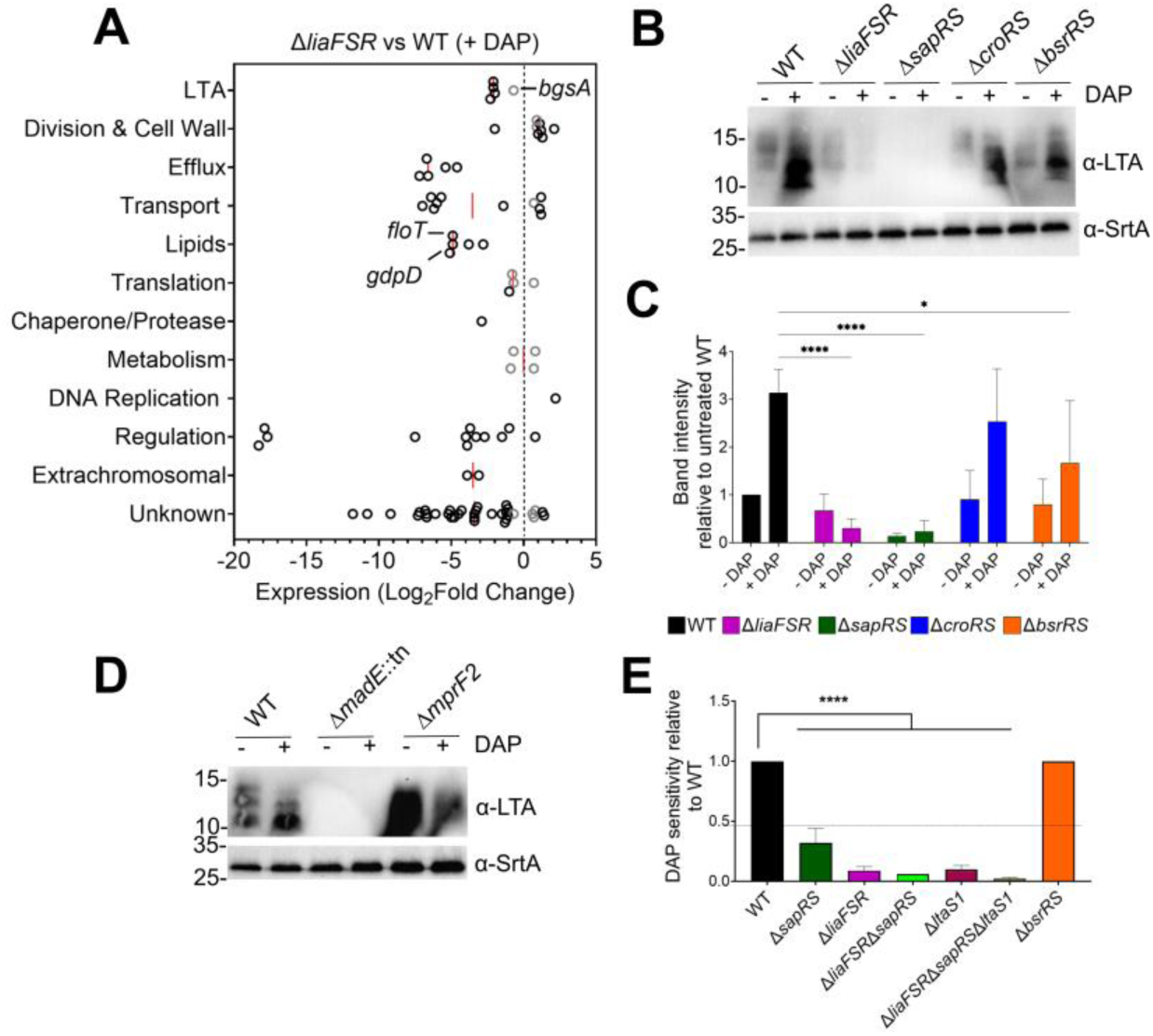
LiaFSR and SapRS control LTA production. (A) RNA-Seq of DAP-treated OG1RF Δ*liaFSR* strain compared to WT cultures. Exponentially grown cells were treated with 2µg/mL DAP for 15 minutes prior to RNA extraction. Analysis was performed on 3 biological replicates per condition. The graph shows the functional classification of differentially expressed genes. Genes exhibiting Log_2_FC ≥ 1.0 with FDR ≤ 0.05 are represented by black circles, while genes exhibiting Log^2^FC ≤ 1.0 with FDR ≤ 0.05 are represented by grey circles. Selected genes are identified by labels. (B) LTA Western Blot of cell extracts of untreated and DAP-treated OG1RF WT, Δ*liaFSR*, Δ*sapRS*, Δ*croRS*, and Δ*bsrRS* strains. SrtA was used as a loading control. Representative blot of three independent biological replicates. Size is denoted in kDa. (C) Band intensity quantification of three independent LTA western blots described in (B) (* P ≤ 0.05, **** P ≤ 0.0001). (D) LTA Western Blot of cell extracts of untreated and DAP-treated OG1RF WT, Δ*mprF2*, and *madE*::tn strains. (E) DAP MIC of OG1RF WT and derivatives in Ca-adjusted BHI media. Bar graph represents DAP sensitivity of strains relative to WT. A 2-fold increase in sensitivity is depicted by a dashed line. Differences analyzed via ANOVA with Tukey’s post-test (**** P ≤ 0.0001).

Although LiaFSR did not control LTA synthesis on the transcriptional level, other TCS including CroRS and BsrRS were induced in response to DAP (**Dataset S1**). CroRS senses cell envelope damage and confers Enterococci with high-level cephalosporin resistance (35). BsrRS (also known as YclRK) is an understudied TCS that transcriptionally responds to cell wall-damaging antibiotics (36–38). Therefore, we performed LTA immunoblots on strains defective for TCS that were induced by DAP, namely Δ*croRS*, Δ*bsrRS*, Δ*liaFSR*, and Δ*sapRS*. Surprisingly, untreated cultures of Δ*sapRS* failed to produce detectable LTA (**Fig. 4B**). When exposed to DAP, LTA levels remained undetectable in Δ*sapRS* and became undetectable in Δ*liaFSR*, indicating that LiaFSR controls LTA production during DAP stress via SapRS. The Δ*bsrRS* strain displayed a two-fold reduction in LTA, while no differences were observed for Δ*croRS* (**Fig. 4C**). Absence of LTA in Δ*sapRS* suggested that LTA production was controlled by an effector of the SapRS system. Possible candidates included MprF2, the main LPG synthase of *E. faecalis*, and MadEFG, a putative efflux pump (12). LTA immunoblots of the Δ*mprF2* strain showed an increase in LTA production even under untreated conditions (**Fig. 4D**). Importantly, no detectable LTA was observed in *madE*::tn, phenocopying the Δ*sapRS* strain. This indicated that LiaFSR and SapRS control LTA synthesis indirectly via transcriptional activation of MadEFG.

### Contribution of LiaFSR and LtaS1 to baseline DAP susceptibility

Inactivation of LiaFSR leads to DAP hyper-susceptibility (39). Since our results indicated that LiaFSR activates LTA synthesis, we next asked whether the hyper-susceptibility of Δ*liaFSR* was due to its inability to elevate LTA and glycolipid synthesis. We determined DAP susceptibility of single (Δ*liaFSR*, Δ*sapRS*, Δ*bsrRS,* Δ*ltaS1),* double (Δ*liaFSR*Δ*sapRS)* and triple (Δ*liaFSR*Δ*sapRS*Δ*ltaS1*) mutant strains (**Fig. 4E**). Though LTA production was 2-fold lower in Δ*bsrRS*, DAP sensitivity was not altered. The Δ*sapRS* strain that failed to produce LTA was ∼4 times more sensitive than the WT. The Δ*liaFSR*, Δ*ltaS1,* and double Δ*liaFSR*Δ*sapRS* strains were highly sensitive to DAP (∼16-fold). However, the triple Δ*liaFSR*Δ*sapRS*Δ*ltaS1* was even more susceptible (∼32-fold). Recently, a mutation in *ltaS* was identified in a clinical DAP^R^ and methicillin^R^ *S. aureus* strain, also revealing LtaS as a contributing factor to high-level DAP^R^ (40). Since *E. faecalis* DAP^R^ strains have a constitutively active LiaFSR and high levels of DGDAG (8, 14–16), we hypothesized that deletion of *ltaS1* would resensitize DAP^R^ strains. However, repeated attempts to delete *ltaS1* in two laboratory-evolved OG1RF DAP^R^ strains harbouring *lia* and *cls* mutations (16) were unsuccessful, suggesting that a gain-of-function in Cls and inactivation of *ltaS1* are synthetic lethal. Nevertheless, taken together, these data suggest that LtaS1 contributes to baseline DAP susceptibility in additional ways, independently of LiaFSR and SapRS.

## Discussion

Our findings reveal that *E. faecalis* mounts a coordinated, phenotypic membrane remodeling response to DAP that mirrors the lipid composition of DAP^R^ strains. Exposure to sub-inhibitory DAP reprograms phospholipid metabolism, diverting phospholipids PG and LPG toward the synthesis of the neutral glycolipid DGDAG and the cell-envelope polymer LTA. This adaptive response is mediated by the two-component signaling network LiaFSR–SapRS–BsrRS and depends on the activity of LtaS1. Together, these systems integrate antibiotic sensing with metabolic flux to fortify the cell envelope against antibiotic stress.

DAP exposure produced a lipid signature nearly identical to that of high-level DAP^R^ strains, characterized by depletion of PG and enrichment of DGDAG. Transcriptomic data indicate that this remodelling is driven both by reduced PG synthesis and by induction of PG-consuming enzymes, including MprF2 and the main LTA synthase LtaS1. Increased activity of these enzymes likely contributes to the net loss of PG and explains accumulation of glycolipids and LTA. Contrary to expectation, PG levels were even lower in the Δ*ltaS1* strain, mimicking findings in Δ*mprF1*Δ*mprF2* and Δ*mprF2*Δ*cls1*Δ*cls2* strains (19, 20), underscoring the remarkable flexibility of *E. faecalis* lipid homeostasis and suggesting compensatory redirection of phospholipid precursors in Δ*ltaS1* through alternative routes such as Cls- or MprF-dependent metabolism.

Pulse-chase labelling experiments revealed that LtaS1 drives *de novo* DGDAG synthesis from newly produced PG while simultaneously using recycled PG and LPG as glycolipid anchors. This dual strategy in PG turnover likely maintains membrane integrity under stress by replenishing neutral lipids while limiting anionic PG pools. This might be especially important in *E. faecalis*, as it lacks the zwitterionic phosphatidylethanolamine (16, 20). The observed LPG recycling parallels the action of the hydrolase AhyD in *E. faecium* (30). In *E. faecalis, ahyD* is co-induced with *mprF2* in response to DAP, which reconciles why LPG hydrolysis is favored during DAP stress even when *mprF2* is induced. PG lysinylation by MprF2 and LTA alanylation by DltABCD modulate surface charge for maintenance of cell membrane function and for antimicrobial protection (41), and both are activated by SapRS (12). Thus, diverting LPG back into PG could satisfy LTA synthesis demands without compromising charge balance.

In *B. subtilis* and *S. aureus,* LtaS localizes at the cell septum where PG is enriched (4, 29, 42). In *E. faecalis,* it is possible that LtaS1 is also located at the septum which is rich in anionic lipid microdomains (PG and CL) (9, 17). The possible synthetic lethal interaction between LtaS1 and Cls in DAP^R^ strains indicates that the activities of these two enzymes need to be carefully balanced to co-exist. We hypothesize that septal LtaS1 location can be advantageous in the context of DAP. The LiaFSR signalling cascade displaces Cls to non-septal locations (9), thereby lowering competition between Cls and LtaS1 to favour LTA and DGDAG synthesis. This would explain why *ltaS1* deletion only affects *de novo* DGDAG synthesis when exposed to DAP.

Our data position LiaFSR as the central regulator that couples membrane damage sensing to adaptive lipid remodelling. LiaFSR activation by DAP stimulates SapRS and its effectors MadEFG, DltABCD, and MprF2-AhyD, thereby inducing LTA synthesis, LTA alanylation, and LPG recycling. The BsrRS system further modulates LTA production, consistent with the genomic linkage between *bsrRS* and *ltaS1*. Although MadEFG has been proposed to be a CAMP efflux pump (12), our data suggest an additional function in LTA biogenesis. One possibility is that MadEFG is a transporter of essential LTA precursors. Alternatively, MadEFG could be involved in c-di-AMP homeostasis. Efflux pumps are associated with c-di-AMP secretion in *L. monocytogenes* during cell wall stress (43), and LTA-defective strains in *S. aureus* compensate loss of LTA via increased intracellular c-di-AMP (44). In *E. faecalis*, the Δ*liaFSR* strain does not produce LTA due to low *madEFG* expression and Δ*liaR* strains have increased intracellular c-di-AMP levels (45). Moreover, strains defective in c-di-AMP metabolism are hyper-sensitive to DAP (46). Through this multi-tiered network, *E. faecalis* likely aligns stress detection with lipid metabolism to preserve cell-surface homeostasis.

We propose a refined model in which LiaFSR signalling initiates a protective shift in membrane composition that precedes and enables the evolution of stable DAP^R^ genotypes (**Fig. 5**). LiaR-mediated displacement of Cls away from the cell septum (9), together with SapRS-dependent activation of MprF2 and MadEFG-dependent LtaS1, redirects lipid flux toward glycolipid accumulation. This remodelling produces a membrane enriched in DGDAG and LTA, structurally reinforcing the cell envelope while creating an environment permissive to subsequent *cls1* gain-of-function mutations. Such a sequence provides a mechanistic explanation for the ordered emergence of *lia* and *cls* mutations observed during the evolution of DAP^R^. Since *cls* mutations confer DAP^R^ (9, 17) and are potentially synthetically lethal to *ltaS1* inactivation, combinatorial therapy using existing LtaS1 inhibitors as adjuvants to DAP is predicted to eliminate the emergence of enterococcal DAP^R^ (47).

**Figure 5.**
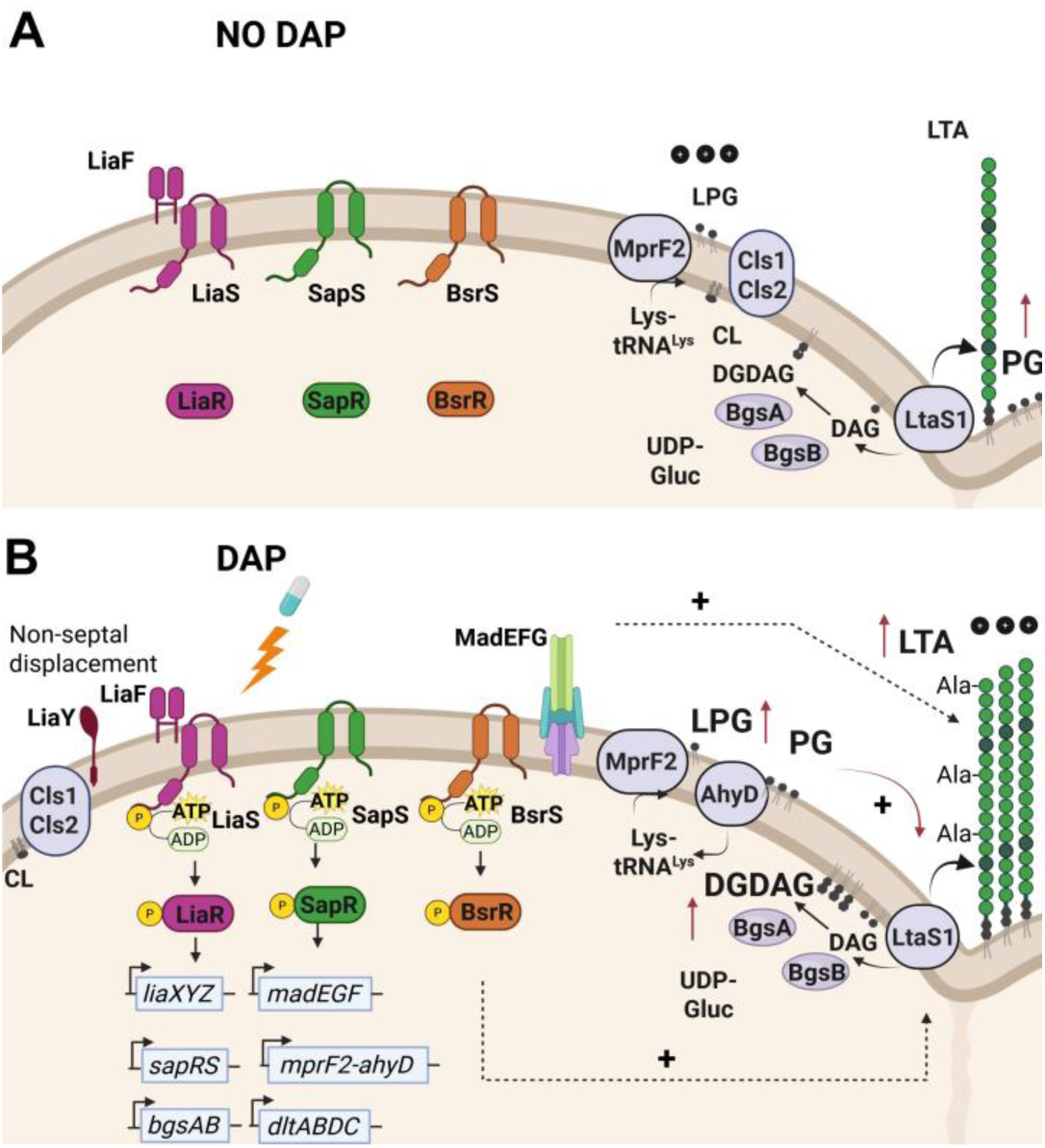
LiaFSR controls adaptive lipid remodelling in response to DAP. (A) In the absence of DAP, the TCS are inactive. The dominant lipid is PG. Cls and LtaS1 are located at the cell septum and produce steady state amounts of CL, DGDAG, and LTA together with BgsAB. MprF2 synthesizes LPG and modulates surface charge. (B) DAP is sensed by LiaX and triggers the LiaFSR signalling cascade that leads to LiaR phosphorylation and transcriptional induction of target genes. On one hand, induction of LiaY displaces Cls to non-septal locations. This displacement decreases PG substrate competition between Cls and LtaS1 and redirects *de novo* PG towards LTA and glycolipid synthesis. Induced transcription of glycosyltransferases *bgsA* and *bgsB* match enhanced LtaS1 activity to support glycolipid synthesis. On the other hand, induction of *sapRS* stimulates the SapRS signalling cascade, which in turn activates transcription of downstream genes. Specifically, MadEFG is required for LTA production, MprF2 and AhyD recycle cationic LPG into LTA-committed DGDAG, while DltABCD alanylates LTA to take over modulation of surface charge. Simultaneous induction of LtaS1 and MprF2 leads to depletion of net PG levels and enrichment of glycolipids. Collectively, LiaFSR controls membrane remodelling in terms of composition and architecture. The resulting membrane configuration enables acquisition of secondary mutations in Cls that lead to high-level DAP^R^.

The glycolipid enrichment in response to DAP is not unique to Enterococci. In fact, higher glycolipid synthesis is associated with DAP^R^ in multiple clinically relevant pathogens including Enterococci, *Clostridioides difficile* and *S. aureus* (14–16, 48–50). Thus, it appears that the combination of lower PG and higher DGDAG fortifies the Gram-positive membrane. The functional consequences of this lipid shift extend beyond DAP tolerance. Replacement of anionic PG with neutral glycolipids may reduce electrostatic attraction to cationic antimicrobials, while accumulation of LTA polymers could increase cell-wall rigidity and protect the septal region from antibiotic attack. This strategy parallels membrane defense in Gram-negative bacteria, where modification of lipid A similarly diminishes cationic drug binding (51). Moreover, comparable lipid transitions occur when *E. faecalis* encounters host-derived fatty acids found in bile and serum (20, 52), suggesting that antibiotic-induced membrane remodeling co-opts an existing environmental adaptation mechanism. By pre-emptively strengthening the membrane in response to its native intestinal niche, Enterococci may achieve both transient protection against a broad spectrum of stressors in the host as well as to exogenous antimicrobials.

In summary, we uncover a unifying mechanism of enterococcal antibiotic resistance in which the LiaFSR–SapRS–BsrRS signalling axis couples antibiotic sensing to dynamic remodelling of membrane lipid composition. LtaS1-dependent redirection of phospholipid flux toward glycolipid and LTA synthesis transiently fortifies the cell envelope and establishes a physiological bridge between short-term tolerance and the subsequent genetic acquisition of resistance. These findings redefine the role of membrane homeostasis in antibiotic adaptation and highlight the bacterial envelope as a highly plastic, actively regulated structure that anticipates rather than merely endures antibiotic stress.

## Materials and Methods

### Bacterial strains and general growth conditions

Bacterial strains and plasmids used in this study can be found in (**Table S1**). *E. faecalis* strains were routinely cultured in Brain Heart Infusion (BHI) media at 37°C (static). *E. coli* strains were grown overnight in LB medium at 37°C in a shaking incubator (175 rpm). When required, antibiotics were added to growth media for stable maintenance of plasmids (**Table S2**). BHI was supplemented with 50 mg/L calcium chloride (CaCl^2^) when cells were treated with DAP to ensure activity (53).

### DAP susceptibility assay

DAP susceptibility of wild-type OG1RF and derivative strains was determined using a modified broth microdilution assay in 96-well plates. Overnight cultures were diluted 1:40 in fresh BHI medium and incubated to early exponential phase (OD_600_ ∼0.25) at 37°C. Erythromycin (10µg/mL) was supplemented to the media for pMSP3535 plasmid-harbouring strains. Then, cells were diluted 1:100 in fresh BHI supplemented with CaCl_2_ to reach a final concentration of ∼10^5^ CFU/mL and were immediately challenged with increasing concentrations of DAP in a 96-well plate. The setup of the 96-well plate follows the standard broth microdilution assay (54). Visual inspection for growth was examined after 18-20h incubation at 37°C. The lowest concentration of DAP that completely inhibited growth was recorded. Independent assays were performed on at least 3 different days.

### Liquid chromatography tandem mass spectrometry (LC-MS/MS) analysis of lipids

Overnight cultures of *E. faecalis* were sub-cultured 1:100 into fresh BHI medium and grown to early exponential phase (OD_600_∼0.25). Then, cultures were treated with 2 µg/mL DAP for 1 hour at 37°C. Cultures were pelleted and washed with 1 mL phosphate buffered saline (PBS) before proceeding with lyophilization.

Lipids were extracted from lyophilized cell pellets using a modified Bligh and Dyer method (**Supporting Information Text**). PG (PG 14:0) and LPG (LPG 16:0) were added to the lyophilized samples to serve as internal standards and lipid extraction was carried out as described previously (16). PG, LPG and DGDAG were quantified by LC-MS/MS using multiple reaction monitoring (MRM) using previously described methodologies (16, 19). For PG and LPG measurements, signal intensities were normalized to the spiked internal standards to obtain relative measurements and further normalized against the initial lyophilized cell pellet weight (16, 19). Due to the lack of commercially available standards for DGDAG and MGDAG, normalization to standards was not performed for DGDAG. Instead, signal intensities were normalized to the weight of the lyophilized cell pellets for semi-quantitative measurements. For data visualization purposes, log_2_-fold changes between treated and untreated samples were calculated and plotted as heatmaps.

### RNA-Sequencing

Overnight cultures were sub-cultured 100-fold into fresh BHI supplemented with 50 mg/L CaCl_2_ and grown at 37°C under static conditions to mid-log phase (OD_600_ ∼0.5). Then, 2 µg/mL DAP was added to the culture and incubated for 15 min. Control cultures were left untreated.

RNA extraction and sequencing was carried out as previously described with minor modifications (**Supporting Information Text)** (21). Cultures were harvested and treated with RNAprotect (Qiagen, USA) to stabilize RNA. Then, cells were lysed, resuspended in TRIzol and chloroform, and the top aqueous phase was mixed with an equal volume of 80% ethanol. Total RNA was isolated using the RNeasy Mini Kit (Qiagen, USA), DNA was depleted via DNase I treatment, and RNA quality was assessed. cDNA library preparation and ribosomal RNA depletion was performed by the in-house sequencing facility using the Illumina Total RNA Prep with Ribo-Zero Plus kit (Illumina, USA). Paired-end sequencing (150×150bp) was performed on the Illumina HiSeqX v2.5 system. Downstream analysis was performed as described in **Supplementary Information Text**.

### Radiolabelled pulse-chase lipid thin-layer chromatography (TLC)

Radiolabeling of lipids was performed as previously described with the following modifications (19, 55). For labelling of total lipids, overnight cultures of *E. faecalis* grown in BHI supplemented with 50 mg/L CaCl_2_ were diluted 1:100 in fresh media spiked with 0.5 µCi/mL [^14^C]-acetate (Perkin Elmer) and grown to OD_600_ of 0.25. Cultures were then treated with 2 µg/mL DAP for 1 hour. For labelling of newly synthesized lipids during DAP exposure, overnight cultures of *E. faecalis* grown in CaCl_2_-supplemented BHI were diluted 1:100 in fresh media and grown to OD_600_ of 0.25. Cultures were then spiked with 0.5 µCi/mL [^14^C]-acetate and treated with 2 µg/mL DAP for 1 hour. For labelling of pre-existing lipids prior to DAP exposure, overnight cultures of *E. faecalis* grown in CaCl_2_-supplemented BHI were diluted 1:100 in fresh media spiked with 0.5 µCi/mL [^14^C]-acetate and grown to OD_600_ of 0.25. Cultures were then pelleted, washed with 1 mL of fresh media, and resuspended in fresh media supplemented with 8.5 mM of unlabelled sodium acetate (Sigma-Aldrich, USA). Cultures were then treated with 2 µg/mL DAP for 1 hour.

For all three setups, after incubation with DAP, lipids were extracted from these cultures and resolved by TLC as previously described (**Supplementary Information Text**) (16, 19). Percentage intensities of each spot relative to the total intensity of all lipid classes were quantified. For data visualization purposes, log_2_-fold changes between treated and untreated samples were calculated and plotted as a heatmap.

### LTA Western Blot

*E. faecalis* strains were grown to early-exponential phase (OD_600_ ∼ 0.25) in BHI supplemented with 50 mg/L CaCl_2_. Cultures were split in two and one tube was treated with 8 µg/ml DAP (Thermo Fisher Scientific, Inc., USA) while the other one was left untreated (control) before continuing incubation at 37°C for 1h. Cells were harvested and resuspended in 1 ml PBS before proceeding with cell lysis and LTA western blotting as previously described with minor modifications (56). Specifically, cell lysis was carried out by 3 cycles of 20s bead beating at room temperature followed by 1 min incubation on ice in between cycles. Primary and secondary antibodies are described in (**Table S2**). To test for restoration of LTA production when *ltaS1* was complemented on the nisin-inducible pMSP3535 plasmid, cell lysates were prepared from *E. faecalis* overnight cultures. Considering leaky expression from the nisin-inducible promoter, cultures were either induced with 25 ng/mL nisin or left untreated to confirm expression.

### Construction of deletion strains

Deletion of *ltaS1*, *ltaS2,* and *sapRS* in the *E*. *faecalis* OG1RF background was carried out using the pGCP213 temperature-sensitive Gram-positive plasmid as previously described and detailed in the **Supplementary Information Text** (57). Construction of Δ*bsrRS* was carried out using the CRISPR-Cas12a system as previously described for *E. faecium* (58). Strains, plasmids, primers and antibiotics used for cloning are described in **Tables S1, S2, and S3**. All gene deletions were confirmed by PCR sequencing of flanking sequences in mutant strains.

### Whole genome sequencing

Oxford Nanopore Technologies was used to perform whole genome sequencing on parent OG1RF and deletion strains Δ*ltaS1*, Δ*ltaS2*, and Δ*ltaS1*Δ*ltaS2* and confirm absence of suppressor mutations. Genomic DNA was extracted from *E. faecalis* WT and mutants using the Wizard HMW DNA Extraction Kit (Promega) following the manufacturer’s instructions. Libraries were prepared using the Rapid Barcoding Kit SQK-RBK114.24 and sequenced in R10.4.1 flow cell on a GridION instrument. Bioinformatic analysis is described in detail in **Supplementary Information Text**.All putative single-nucleotide polymorphisms (SNPs) and insertions/deletions (indels) flagged by the pipeline were manually verified by read visualization using the Integrative Genomics Viewer (IGV)

### Construction of complementation strains

Complementation of the Δ*ltaS1* strain was performed using the nisin-inducible expression vector pMSP3535 (59). The coding sequence of *ltaS1*, spanning the entire gene and including a predicted rho-independent terminator region, was amplified from genomic *E. faecalis* OG1RF DNA using the ltaS1-comp primers listed in **Table S3**. Then, vector pMSP3535 and insert were digested with BamHI and XbaI and ligated using T4 ligase to yield the plasmid pMSP-*ltaS1*. The construct was transformed into *E. coli* Stellar cells for propagation, and transformants were screened with pMSP3535 screening primers to isolate positive clones and confirmed by Sanger sequencing. Then, pMSP3535 (empty plasmid) and pMSP-*ltaS1* were electroporated into OG1RF wild-type or Δ*ltaS1* strains using standard protocols (60) and presence of the plasmid was confirmed by PCR screening with pMSP3535 screening primers.

### Statistical analysis

Data sets were analyzed using GraphPad Prism 10 software. Differences in LC-MS/MS-detected lipid classes between treated and untreated OG1RF WT cells were determined using Students *t* test. Transcriptional differences between DAP-treated and untreated cells (RNA-Seq) were assessed using edgeR (version 3.42.4) and setting an FDR-adjusted *p*-value of 0.05 as cutoff. Differences in LC-MS/MS-detected lipid species, TLC-resolved lipid classes, western blot band quantification, and DAP sensitivity were determined using ANOVA followed by a multiple comparison test as specified in corresponding figure legends.

## Data availability

All RNA-sequencing datasets have been deposited in the NCBI Gene Expression Omnibus (GEO) database under accession number GSE309958.

## Acknowledgments

We extend our appreciation to James Collins from the University of Louisville for providing the pJC005.em plasmid (Addgene plasmid # 182738). We thank our colleagues at the SCELSE Sequencing Facility for performing library preparation and RNA sequencing. We thank Chng Shu-Sin, and the members from his lab, Tan Wee Boon and Chen Yushu, for their assistance with radiolabeling of lipids and TLC. We also appreciate the bioinformatic support of Fedor Bezrukov and Julien Prados of the Bioinformatics Support Platform at the University of Geneva.

This work was supported by funding to K.A.K. from the Société Académique de Genève and from the Swiss National Science Foundation grant number 310030_219227. This work was partially supported by the Singapore Centre for Environmental Life Sciences Engineering (SCELSE), funded by the National Research Foundation and Ministry of Education, Singapore under its Research Centre of Excellence Program, as well as by the Singapore Ministry of Education under its Tier 1 program (2020-T1-002-103) and the National Medical Research Council Open Fund (MOH-000645), both awarded to K.A.K.. Z.J.N. and this work was also partially supported by the National Research Foundation, Singapore, under its Campus for Research Excellence and Technological Enterprise (CREATE) program, through core funding of the Singapore-MIT Alliance for Research and Technology (SMART) Centre, Antimicrobial Resistance Interdisciplinary Research Group (AMR IRG).

