## Supporting Information for "A two-component system signaling hub controls enterococcal membrane remodeling in response to daptomycin"

**Supporting Information Text**

**Extended methods**

**Liquid chromatography tandem mass spectrometry (LC-MS/MS) analysis of lipids**

Overnight cultures of *E. faecalis* were sub-cultured 1:100 into fresh BHI medium and grown to early exponential phase ($OD_{600}$~0.25). Then, cultures were treated with 2 μg/mL DAP for 1 hour at 37°C. Cultures were pelleted and washed with 1 mL phosphate buffered saline (PBS) before proceeding with lyophilization.

Lipids were extracted from lyophilized cell pellets using a modified Bligh and Dyer method, where the extraction solvent contained chloroform/methanol in a ratio of 1:2 (vol/vol) as previously described (1, 2). PG (PG 14:0) and LPG (LPG 16:0) were prepared as 0.1 mg/mL stocks in the extraction solvent and 45 μL of PG 14:0 and 36 μL of Lys-PG 16:0 were added to the lyophilized samples to serve as internal standards. Additional extraction solvent was then added to a final volume of 900 μL and lipid extraction was carried out as described previously (1). The dried lipid extract was resuspended in 30 μL chloroform/methanol (1:1 vol/vol) and stored at −80°C until the mass spectrometry analysis was performed.

PG, LPG and DGDAG were quantified by LC-MS/MS using multiple reaction monitoring (MRM) using previously described methodologies (1, 2). An Agilent 6490 QqQ mass spectrometer connected to a 1290 series chromatographic system was used with a Kinetex 2.6 μm HILIC column (100 Å, 150 × 2.1 mm) (Phenomenex, USA). Electrospray ionization (ESI) was used to ionize lipids. Each lipid molecular species was analysed using a targeted MRM approach containing transitions for known precursor/product mass-to-charge ratio (m1/m3). For PG and LPG measurements, signal intensities were normalized to the spiked internal standards to obtain relative measurements and further normalized against the initial lyophilized cell pellet weight (1, 2). Due to the lack of commercially available standards for DGDAG and MGDAG, normalization to standards was not performed for DGDAG. Instead, signal intensities were normalized to the weight of the lyophilized cell pellets for semi-quantitative measurements. For data visualization purposes, $\log_2$-fold changes between treated and untreated samples were calculated and plotted as heatmaps.

**RNA-Sequencing**

Overnight cultures were sub-cultured 100-fold into fresh BHI supplemented with 50 mg/L $CaCl_2$ and grown at 37°C under static conditions to mid-log phase ($OD_{600}$ ~0.5). Then, 2 μg/mL DAP was added to the culture and incubated for 15 min. Control cultures were left untreated. For RNA extraction, cultures were harvested and treated with RNAprotect (Qiagen, USA) to stabilize RNA. Then, cells were lysed with 20 mg/mL lysozyme (Sigma-Aldrich, USA) in lysis buffer (10 mM Tris-HCl at pH 8, 50 mM NaCl, 1 mM EDTA, 0.75 M Sucrose) at 37°C for 1 h. Following centrifugation, the pellet was resuspended in 1 mL of TRIzol and 200 μL of chloroform, and incubated at 4°C for 2 min. After centrifuging at 12,000g at 4°C for 15 min, the top aqueous phase was mixed thoroughly with an equal volume of 80% ethanol. Total RNA was then isolated using the RNeasy Mini Kit

(Qiagen, USA) according to the manufacturer's recommendations. DNA was depleted via on-column DNase I treatment (Qiagen, USA). RNA quality (RINe ≥8) was assessed using the RNA ScreenTape system (Agilent Technologies, USA), while DNA contamination (≤10%) was quantified using the Qubit RNA BR and dsDNA HS Assay kits (Invitrogen, USA). cDNA library preparation and ribosomal RNA depletion was performed by the in-house sequencing facility using the Illumina Total RNA Prep with Ribo-Zero Plus kit (Illumina, USA). Paired-end sequencing (150x150bp) was performed on the Illumina HiSeqX v2.5 system. The quality of the sequencing data was assessed using FastQC (version 0.11.9)(3). Adapter and quality trimming were performed using bbduk (BBMap version 38.96)(4). The trimmed reads were mapped to the *E. faecalis* OG1RF reference genome (GenBank accension CP002621.1) using the BWA-MEM algorithm of Burrows-Wheeler Aligner (version 0.7.17-r1198-dirty)(5) with the default settings. Transcript abundance was determined using htseq-count (HTSeq version 2.0.1) (3) with intersection-strict mode. Differential gene expression (DGE) analysis was performed in R (version 4.3.1) using edgeR (version 3.42.4)(6).

**Radiolabelled pulse-chase lipid thin-layer chromatography (TLC)**

Radiolabeling of lipids was performed as previously described with the following modifications (2, 7). For labelling of total lipids, overnight cultures of *E. faecalis* grown in BHI supplemented with 50 mg/L $CaCl_2$ were diluted 1:100 in fresh media spiked with 0.5 µCi/mL [$^{14}$C]-acetate (Perkin Elmer) and grown to $OD_{600}$ of 0.25. Cultures were then treated with 2 µg/mL DAP for 1 hour. For labelling of newly synthesized lipids during DAP exposure, overnight cultures of *E. faecalis* grown in $CaCl_2$-supplemented BHI were diluted 1:100 in fresh media and grown to $OD_{600}$ of 0.25. Cultures were then spiked with 0.5 µCi/mL [$^{14}$C]-acetate and treated with 2 µg/mL DAP for 1 hour. For labelling of pre-existing lipids prior to DAP exposure, overnight cultures of *E. faecalis* grown in $CaCl_2$-supplemented BHI were diluted 1:100 in fresh media spiked with 0.5 µCi/mL [$^{14}$C]-acetate and grown to $OD_{600}$ of 0.25. Cultures were then pelleted, washed with 1 mL of fresh media, and resuspended in fresh media supplemented with 8.5 mM of unlabelled sodium acetate (Sigma-Aldrich, USA). Cultures were then treated with 2 µg/mL DAP for 1 hour.

For all three setups, after incubation with DAP, lipids were extracted from these cultures as previously described and resuspended in 50 µL of chloroform-methanol solution (1:1 vol/vol) (1). Radioactive counts of the lipid extracts were measured, and the lipid extracts spotted on to silica gel-coated thin-layer chromatography (TLC) plates (Merck, USA) while normalizing to scintillation counts as previously described (2). TLC plates were developed in preequilibrated TLC chambers with a chloroform:methanol:water (65:25:4) solvent system. TLC plates were then visualized by exposure to a storage phosphor screen (GE health care, USA) overnight, and read using a Storm Phosphorimager (GE health care, USA). For data visualization purposes, $log_2$-fold changes between treated and untreated samples were calculated and plotted as a heatmap.

**Construction of deletion strains**

Deletion of *ltaS1*, *ltaS2,* and *sapRS* in the *E. faecalis* OG1RF background was carried out using the pGCP213 temperature-sensitive Gram-positive plasmid as previously described (8). Construction of Δ*bsrRS* was carried out using the CRISPR-Cas12a system as previously described for *E. faecium* (9). Strains, plasmids, primers and antibiotics used for cloning are described in **Tables S1, S2, and S3**. All gene deletions were confirmed by PCR sequencing of flanking sequences in mutant strains.

Briefly, ~ 0.5 kb PCR products flanking the *sapRS, ltaS1,* or *ltaS2* coding sequences were amplified with the primers listed in (**Table S3**). Restriction enzyme sites were created as silent mutations whenever possible to aid in cloning. The amplicons included the first and last ~15 amino acid codons of the coding DNA to avoid unanticipated polar effects. The flanking regions were fused together by overlap extension PCR to generate the final insert. Then, insert and pGCP213 were digested with appropriate restriction enzymes and ligated with T4 ligase following the manufacturer's instructions. The generated plasmids were transformed into competent *E. coli* Stellar cells, transformants were screened with M13 primers and verified by PCR and by Sanger sequencing. Electroporation into electrocompetent *E. faecalis* OG1RF and final isolation of mutant strains were carried out as previously described (10). Colonies were screened with the respective screening primers to identify clones harboring the expected gene deletions. Double mutants were obtained electroporating the different pGCP213 constructs into electrocompetent Δ*ltaS2* or Δ*liaFSR single mutants.* A triple mutant was obtained by electroporating the pGCP-*ltaS1* plasmid into the Δ*liaFSR*Δ*sapRS* double mutant strain.

For construction of Δ*bsrRS*, a 1.3 kb synthetic insert was designed, composed of: (i) the Cas12a target sequence, located immediately downstream of a TTTV protospacer adjacent motif (PAM) within the gene of interest, (ii) ~500 bp sequences homologous to the regions upstream and downstream of the target gene for recombination. The sequence of the synthetic insert is provided in **Table S3**. The synthetic insert was amplified using primers insert-F and insert-R and subsequently cloned into the plasmid pJC005, which had been linearized by PCR with the primers pJC005-inv-1 and pJC005-inv-2, using in vivo assembly in *E. coli* Stellar competent cells (11, 12). The resulting recombinant plasmid was extracted, verified by Sanger sequencing (Microsynth), and electroporated into competent *E. faecalis* OG1RF prepared by overnight growth in BHI 7% glycine, 0.5M sucrose. Transformants were selected on Brain Heart Infusion (BHI) agar supplemented with 50 µg/mL erythromycin. Cas12a expression was induced with 250 ng/mL anhydrotetracycline (ahTC) in the presence of 50 µg/mL erythromycin. Successful gene knockout was confirmed by PCR using primers screen-F and screen-R. Plasmid curing was carried out by two successive passages in BHI broth containing ahTC but lacking erythromycin.

**Whole genome sequencing**

Oxford Nanopore Technologies was used to perform whole genome sequencing on parent OG1RF and deletion strains Δ*ltaS1*, Δ*ltaS2*, and Δ*ltaS1*Δ*ltaS2* and confirm absence of suppressor mutations. Genomic DNA was extracted from *E. faecalis* WT and mutants using the Wizard HMW DNA Extraction Kit (Promega) following the manufacturer's instructions. Libraries were prepared using the Rapid Barcoding Kit SQK-RBK114.24 and sequenced in R10.4.1 flow cell on a GridION instrument. Reads were basecalled with Dorado (v.0.9.2) in super accuracy mode, genomes were assembled with flye (v.2.9.1) and polished with medaka (v.2.0.1). To refine SNPs and indel detection reads, knockout strains were directly compared with WT reads and with the *E. faecalis* OG1RF reference genome (NCBI assembly accession: GCA_000172575.2) using the medaka pipeline available at https://github.com/BioinfoSupport/nanopore_compare. All putative single-nucleotide polymorphisms (SNPs) and insertions/deletions (indels) flagged by the pipeline were manually verified by read visualization using the Integrative Genomics Viewer (IGV) (13).

### Figures

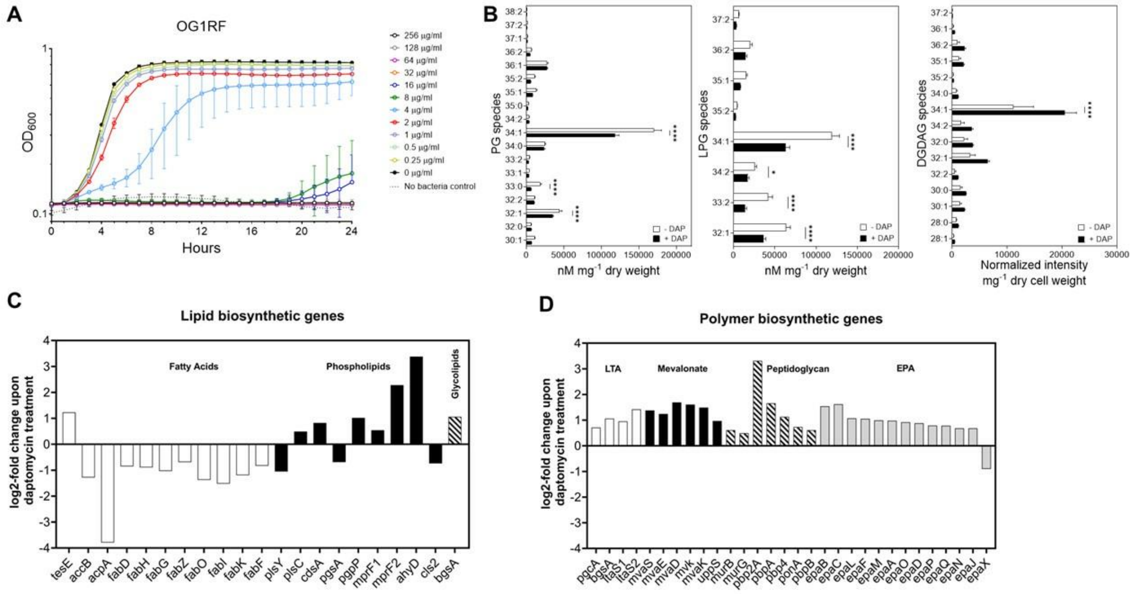


**Fig. S1**. (A) Growth of WT *E. faecalis* OG1RF in BHI supplemented with $CaCl_2$ and increasing concentrations of DAP. Growth was monitored by measuring $OD_{600}$ every hour for up to 24h using a Tecan plate reader. The graph shows the average and standard deviation of three biological replicates. Independent experiments were performed twice with equal results. (B) LC-MS/MS quantification of individual PG, LPG, and DGDAG species in treated and untreated WT *E. faecalis* OG1RF. Carbon length and saturation status of fatty acids are shown in the y-axis. Each bar represents the mean ± standard error of 3 to 5 biological replicates. Differences in lipid content were analysed via a two-way ANOVA followed by Šídák's post-test (* P ≤ 0.05, **** P ≤ 0.0001). (C, D) Transcriptional expression of selected (B) lipid and (C) cell envelope-associated polymer biosynthetic genes during DAP treatment. The *E. faecalis* OG1RF strain was treated with 2µg/mL DAP for 15 min prior to RNA extraction and downstream RNA-Seq analysis. The bar graphs show the mean $log_2$ fold-change of three biological replicates comparing treated to untreated OG1RF cultures. All values shown were statistically significant (FDR < 0.05). LTA = Lipoteichoic Acid, EPA = Enterococcal Polysaccharide Antigen.

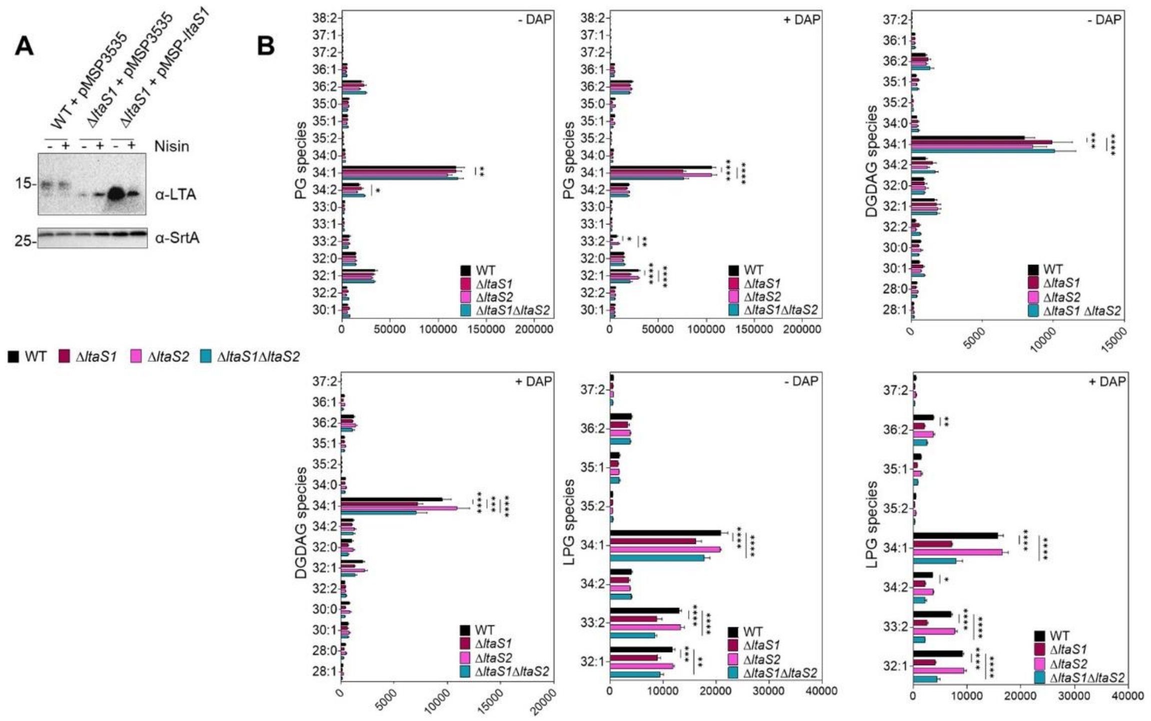


**Fig. S2**. (A) LTA Western Blot of cell extracts of OG1RF WT and Δ*ltaS1* carrying the empty pMSP3535 plasmid or Δ*ltaS1* carrying the pMSP-*ltaS1* vector. Overnight cultures were incubated with or without 25 ng/mL nisin prior to cell lysis to determine LTA production under induced and uninduced conditions, respectively. SrtA was used as a loading control. Size is denoted in kDa. Of note, the nisin-inducible promoter was reported to be leaky (14).(B, C, D) Densitometric analysis of [14C] radiolabeled 1D TLC spots of (B) total lipids, (C) newly synthetized lipids, and (D) pre-existing lipids in parent and LTA-defective strains in the absence of DAP (-DAP). Each bar represents the mean ± standard error of measurement calculated from 2 biological replicates. Spot identities were determined using lipid standards. Results were analyzed via two-way ANOVA. Differences were not statistically significant. (E, F, G) LC-MS/MS quantification of total (E) PG, (F) LPG, and (G) DGDAG content of untreated WT and LTA-deficient *E. faecalis* cells. PG and LPG were normalized to internal standard and cell pellet weight, while DGDAG intensity was normalized to cell pellet weight for a semi-quantitative measurement. Each bar represents the mean ± standard error of 3 to 5 biological replicates. Differences in lipid content were analysed via one-way ANOVA. (* P ≤ 0.05, ** P ≤ 0.01).

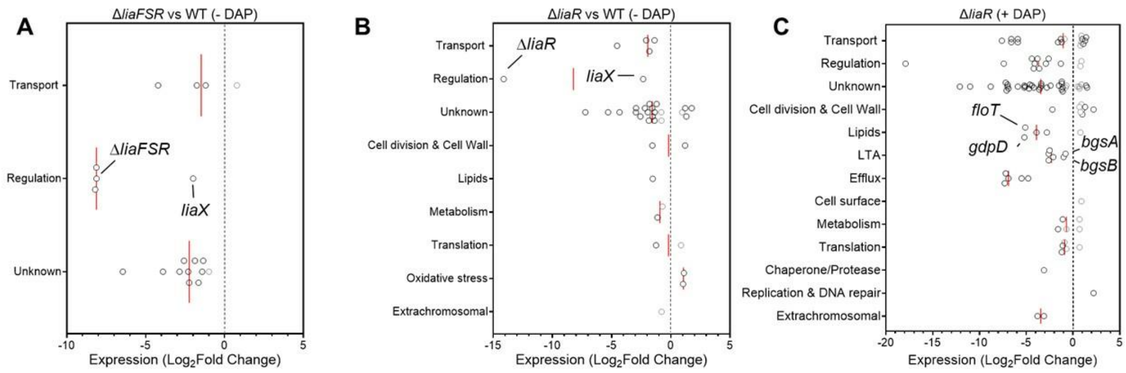


**Fig. S3.** (A-C) RNA-Seq of (A) untreated OG1RF Δ*liaFSR*, (B) untreated OG1RF Δ*liaR*, and (C) DAP-treated OG1RF Δ*liaR* compared to their respective WT controls. Exponentially grown cells were either treated with 2μg/mL DAP or left untreated for 15 minutes prior to RNA extraction. Analysis was performed on 3 biological replicates per condition. The graph shows the functional classification of differentially expressed genes. Genes exhibiting Log2FC ≥ 1.0 with FDR ≤ 0.05 are represented with black circles, while genes exhibiting Log2FC ≤ 1.0 with FDR ≤ 0.05 are represented with grey circles; selected genes are labelled for ease of identification.

**Tables**

**Table S1. Bacterial strains and plasmids**

| Strain | Description[a] | Reference |
|---|---|---|
| *E. faecalis* | | |
| OG1RF | Reference sequenced strain; Fus[R] and Rif[R] | (15, 16) |
| ∆*ltaS1* | OG1RF *ltaS1* deletion | This work |
| ∆*ltaS2* | OG1RF *ltaS2* deletion | This work |
| ∆*ltaS1*∆*ltaS2* | OG1RF *ltaS1* and *ltaS2* deletion | This work |
| OG1RF + pMSP3535 | Parent strain carrying the empty pMSP3535 plasmid | This work |
| ∆*ltaS1* + pMSP3535 | ∆*ltaS1* strain carrying the empty pMSP3535 plasmid | This work |
| ∆*ltaS1* + pMSP-*ltaS1* | ∆*ltaS1* strain carrying the pMSP-*ltaS1* construct | This work |
| ∆*liaFSR* | OG1RF *liaFSR* deletion | (8) |
| ∆*liaR* | OG1RF *liaR* deletion | (8) |
| ∆*sapRS* | OG1RF *sapRS* deletion | This work |
| ∆*croRS* | OG1RF *croS* deletion, no expression of *croR* | (17) |
| ∆*bsrRS* | OG1RF *bsrRS* deletion | This work |
| ∆*mprF2* | OG1RF *mprF2* deletion | (18) |
| *madE::tn* | Tn insertion at site 2392085, Cm[R] | (19) |
| ∆*liaFSR*∆*sapRS* | OG1RF *liaFSR* and *sapRS* deletion | This work |
| ∆*liaFSR*∆*sapRS*∆*ltaS1* | OG1RF *liaFSR, sapRS* and *ltaS1* deletion | This work |
| *E. coli* | | |
| Stellar | Host strain of pGCP213, pMSP3535, and pJC005 | Lab stock |
| DH5α | Host strain of empty pMSP3535 plasmid | Lab stock |

| Plasmid | Description[a] | Reference |
|---|---|---|
| pGCP213 | Temperature-sensitive plasmid used for allelic exchange; Erm[R]: 500 µg/mL *E. coli;* 25 µg/mL *E. faecalis* | (10) |
| pGCP-*ltaS1* | pGCP213 carrying *ltaS1* deletion insert; Erm[R] | This work |
| pGCP-*ltaS2* | pGCP213 carrying *ltaS2* deletion insert; Erm[R] | This work |
| pGCP-*sapRS* | pGCP213 carrying *sapRS* deletion insert; Erm[R] | This work |
| pJC005.em | CRISPR-Cas12a genome editing plasmid; Erm[R]: 200 µg/mL *E. coli;* 50 µg/mL *E. faecalis* | (9) |
| pJC-*bsrRS* | pjC005 carrying *bsrRS* deletion insert; Erm[R] | This work |
| pMSP3535 | Gram-positive, nisin-inducible expression vector; Erm[R]: 300 µg/mL *E. coli,* 10 µg/mL *E. faecalis* | (14) |

| | | |
|---|---|---|
| pMSP-*ltaS1* | pMSP3535 carrying the *ltaS1* gene under the control of the nisin-inducible promoter; $Erm^R$ | This work |

[a]Fus: fusidic acid; Rif: rifampicin; Erm: erythromycin

**Table S2. Antibodies and antibiotics used in this study.**

| Antigen | Size (kDa) | Primary antibody (concentration; host) | Secondary antibody (concentration; host) | Reference |
|---|---|---|---|---|
| LTA | 10-14 | Hycult Clone 55 (1:1000; Mouse) | α-mouse IgG HRP (1:5000; Goat) | (2) |
| SrtA | 27.1 | α-SrtA (1:250; Mouse) | α-mouse IgG HRP (1:1000; Goat) | (20) |

| Antibiotic | Vendor | Reference | Stock concentration |
|---|---|---|---|
| Daptomycin | Thermo Scientific | 461375000 | 1 mg/mL |
| Erythromycin | Thermo Scientific | 227330050 | 100 μg/mL |
| Nisin | Sigma Aldrich | N5764-5G | 1 μg/mL |
| Anhydrotetracycline | Thermo Scientific | 24120625 | 1 mg/mL |

**Table S3. Primers used in this study.**

| Primers | Sequence* | Application |
|---|---|---|
| M13 F | 5’GTAAAACGACGGCCAGT3’ | pGCP213 |
| M13 R | 5’CAGGAAACAGCTATGAC3’ | screening |
| yxdJK_del1 | 5’CTCAAATCCAGCTCGAGCAAATGAAG3’ | |
| yxdJK_del2 | 5’ATAATAACTGAGGGAAGGCTTCGACAATCATGATTTTAGCC3’ | |
| yxdJK_del3 | 5’AAATCATGATTGTCGAAGCCTTCCCTCAGTTATTATAATGAAG3’ | *sapRS* |
| yxdJK_del4 | 5’CAACGCCCGGATCCCTTTATTAAAATC3’ | deletion |
| yxd screen F | 5’GCCGATTTTTTTATAAGATTATGC3’ | |
| yxd screen F | 5’GTTGATTCTGTCCCAAACTCC3’ | |
| ltaS1_del1 | CCGCTTCTCTCTAGATATTTCAAAGAG | |
| ltaS1_del2 | CCCGTTTTTTGATAGTTTTCAAAATAAACGTCC | |
| ltaS1_del3 | GGACGTTTATTTTGAAAACTATCAAAAAACGGG | *ltaS1* |
| ltaS1_del4 | GAGACCACTGCAGAATAGCATCAC | deletion |
| ltaS1 screen F | CTTGAAGTTTCATTCTGTACACC | |
| ltaS1 screen R | TCTGGCATCGTTCGTAAAAAC | |
| ltaS2_del1 | GCTATATCTGCAGGAGATGAGAGGC | |
| ltaS2_del2 | AGTCGGATAATTTTGGACTTAAAAAGAGTCCTAATCGGTT | |
| ltaS2_del3 | AACCGATTAGGACTCTTTTTAAGTCCAAAATTATCCGACT | *ltaS2* |
| ltaS2_del4 | CTTTTTTGCTAGATAAGCTTAGCCCATCATCAA | deletion |
| ltaS2 screen F | TGCGGTGACATGTAAGCAA | |
| ltaS2 screen R | GCCTGTTGTATTTTCTGTTGATGG | |
| pJC005-inv-1 | CTGGCATCTTTTTATTTAGGGATTTCTCAC | pJC005 |
| pJC005-inv-2 | GACATGCAAGCTTGGCACTGGC | cloning |
| insert-F | GTGAGAAATCCCTAAATAAAAAGATGC | |
| insert-R | GCCAGTGCCAAGCTTGCATGTC | *bsrRS* deletion |
| yclRK-screen-F | GGTCAATTCAGGAAACCGC | |

| | | |
|---|---|---|
| yclRK-screen-R | CTCTTGTTTGCGGGACTCC | |
| ltaS1-comp-F | AATCATGGATCCGAAAGGACGTTTATTTTG | |
| ltaS1-comp-R | TCTCATTCTAGATCACCTCTCTAAACTATCTTTAG | *ltaS1* |
| pMSP screen F | ACCCGGTTGTAAAACAGGAG | complement |
| pMSP screen R | ACGACTCACTATAGGGAGACC | |

*Underlined bases correspond to restriction sites included to clone PCR products.

Synthetic inserts

>yclRK-synthetic-insert
GTGAGAAATCCCTAAATAAAAAGATGCCAGTGTGCTGGAATTCGTCCACTAGTCATTCTACA
GTTTATTCTTGACATTGCACTGTCCCCCTGGTATAATAACTAT**AATTTCTACTCTTGTAGAT**ct
cttccgtcatgccgattcctg**AATTTCTACTCTTGTAGAT**aaaaatacggcgctcattgtaaaaagccaattcatgaccac
gctgtttgacatgttcatgcagtgaaacacattcctctgttgtaaattcatctgaataaatcgtttttccttcataatgaataaattgcccgttcatt
acaatcgcagaatcaatgccactgtctttcataattggctgaatctcacacaaggtccgtcctgtcgcaattaaaggtaaaatttgattgtctt
tcaaggccgccactgctgcggtaatctcaggagttatttgcgatgttccatctaacaaagttccatctaaatcaaaaaaagtaatagctttc
attttatccataccctaacatcctctctacgcttgttattttaacaaacttttctgaaaagacaatgggttcctttcgttttctttaaaagagcagga
aaaaaagtgatttttgatggttctgtaaaggatttttgaagaaatatagaatcggttggtacaaattcaattttagagtatactacaatagagc
agttttatgaaaagtgaataaggagtgaatgaaagaaaacgtctggtcacaatcagtgccagacgtttttttagtttaataacgcatgtaga
gacaaggggcgcacatagggactatcaataatttgtggatcctttgtcagaaatagcaagcgcttgtttctttttgaactgctatttttaattga
ggtaaaatttcttgacgttgtccaatcgttaatgaagaaaagatatcatcaatcactaacgtacttgtttttgaaaacaaaagttgtattatgcg
cattttaatttttcaaaagtcgtaaaagaagtaaaggaacgaccaagtaaggcaggttctaattggaacacctggaaagcttcggctag
aactacttttcgatcccgttctttgatcgtactagcgataaataaattttcttctaatggcaagtagggcaaaaaggtttcactagaagaaag
cctagtaatggttagttggtttgtttggaaaaaatcggccaatgcgagttcttgttctggagcagaataaatgattccataaacttctcgcaaa
gctaacgcgtcgaatgaggatttGACATGCAAGCTTGGCACTGGC

**Dataset S1 (separate file).** RNA-Seq results comparing daptomycin-treated vs untreated wild-type *E. faecalis* OG1RF. Genes with significant differences in mRNA abundance were determined using a P value and false discovery rate (FDR) cut-off of 0.05. The $log_2$FC represents the $log_2$ fold change between biological triplicates of exponentially growing OG1RF when treated for 15 min with 2μg/mL of daptomycin in BHI supplemented with 50mg/L $CaCl_2$. Functional classification of genes was manually curated.

**Dataset S2 (separate file).** RNA-Seq results comparing treated or untreated OG1RF Δ*liaFSR* and Δ*liaR* strains to treated or untreated wild-type *E. faecalis* OG1RF strain, respectively. Genes with significant differences in mRNA abundance were determined using a P value and false discovery rate (FDR) cut-off of 0.05. The $log_2$FC represents the $log_2$ fold change between biological triplicates of exponentially growing OG1RF in BHI supplemented with 50mg/L $CaCl_2$. Functional classification of genes was manually curated.
